# A set of constitutive promoters with graded strengths for gene expression in diverse cyanobacterial strains

**DOI:** 10.64898/2026.03.25.714268

**Authors:** Kevin P Trieu, Bryan Bishé, Arnaud Taton, Brian P. Tieu, James W. Golden

## Abstract

Cyanobacteria have garnered interest as promising biological platforms for producing renewable biofuel, chemical feedstock, and bioactive molecules. For biotechnology applications, robust well-characterized genetic tools are required for genetically modifying cyanobacteria, but these tools are often developed for specific model strains. Here, we used broad host-range RSF1010-based plasmids to characterize a set of orthogonal constitutive promoters in diverse cyanobacterial strains. The promoters are random variants of the synthetic *Escherichia coli* PconII promoter. A library of PconII promoters driving a fluorescent reporter gene was first evaluated in *Synechococcus elongatus* and found to have a wide range of gene expression levels. A set of 25 promoter variants with graded strengths was selected after characterization in *S. elongatus* and three additional model cyanobacterial strains. To demonstrate the utility of these promoters, we isolated new genetically tractable cyanobacterial strains with high salt and alkalinity tolerance and transferred the subset of promoters into one of these newly isolated strains. Similar to the results with model strains, the subset of promoters had a wide range of expression levels in the non-model strain. These characterized promoters expand the genetic tools available for genetic engineering of model and non-model cyanobacterial strains.

**Importance:** The use of cyanobacteria to produce renewable products will require engineered expression of many genes that affect cell growth, metabolism, and agronomic properties, leading to efficient production of biomass and desired products. Engineering the strength of gene transcription is an important element of overall gene expression levels. The set of constitutive promoters described here, with a wide range of expression strengths characterized in several diverse cyanobacterial strains, provides an important resource for genetic engineering required for biotechnology applications.

**Research Areas:** Microbial genetics, plasmids and other genetic constructs, biotechnology

**Journal Secction:** Biotechnology

## Introduction

Genetic tools are fundamental to basic research and biotechnology for manipulating gene expression and metabolic pathways. Most available tools for cyanobacteria have been developed to study fundamental cell biology processes in particular species, especially in model strains that grow well in the laboratory and are amenable to genetic manipulations (1–4). While such tools have been used and sometimes further developed for engineering cyanobacteria to produce biofuels and other biomolecules (5, 6), yields remain low, particularly, in comparison to those obtained with heterotrophs like *Escherichia coli* (hereafter *E. coli*) (7). To advance sustainable production of biochemicals and other biotechnological applications using cyanobacteria, additional genetic tools and strains are needed. Considering the diversity of the cyanobacterial phylum, some genetic tools may not be compatible across different strains, especially non-model strains (1). Types of improvements needed to optimize cyanobacterial growth and production of desired products include minimizing competing native metabolic pathways, improving metabolic flux into targeted pathways, and increasing cell tolerance to high concentrations of biochemical end products (8). The success of these modifications in increasing production of desired compounds and limiting production of harmful byproducts will require robust methods for controlling gene expression levels.

Heterologous promoters, notably those from *E. coli,* have been adapted for use in cyanobacteria. For example, the inducible *E. coli* Ptrc promoter has been used in *Synechocystis* PCC 6803 (hereafter *Synechocystis*), *Synechococcus elongatus* PCC 7942 (hereafter *S. elongatus*), and *Anabaena* PCC 7120 (hereafter *Anabaena*) (9–11). However, Ptrc transcriptional control is imperfect in cyanobacteria, with a level of leaky basal expression occurring in the absence of inducer (12). Other *E. coli* promoters tested in cyanobacteria include Plac and Ptet; however, both exhibit low activity in *Synechocystis* (10). Other engineering strategies, such as cyanobacterial-specific promoter libraries, have been used to yield a wider range of promoter expression levels. Markley *et al.* mutagenized a truncated sequence of the strong *cpcB* promoter from *Synechococcus* PCC 7002, generating a set of 11 promoters spanning three orders of magnitude of YFP expression levels (13). They also tested a set of *E. coli* promoters from the BioBrick promoter family, which yielded an expression range of 2.5 orders of magnitude. Notably, there was only a weak correlation between *E. coli* and *Synechococcus* PCC 7002 expression levels measured by reporter fluorescence. This discrepancy could be attributed in part to differences in the promoter consensus sequences between the two organisms (13). These results illustrate that *E. coli* promoter libraries may not function as expected and must be empirically tested in the cyanobacterial strains in which they will be used. Furthermore, even within the cyanobacterial phylum, promoter sequences are not well conserved and there is significant evolutionary divergence of sigma factors, which suggests that promoters may not function equally well in different cyanobacterial strains (14, 15).

In *E. coli,* the sigma70 housekeeping sigma factor binds to -10 and -35 consensus regions upstream of the transcription start site to recruit RNA polymerase and initiate transcription (16). In *E. coli*, the -10 region is conserved across sigma70 promoters, with a consensus sequence ‘TATAAT’ between positions -12 and -7 relative to the transcriptional start site (16). The positions -15 and -14 of sigma70 promoters are partially conserved, comprising the extended -10 region. These conserved regions influence promoter strength, which can be modulated by altering the nucleotide bases in those regions (17).

Among *E. coli* promoters, the synthetic conII promoter (PconII) has been used for constitutive gene expression in different cyanobacterial genera, including strains of *S. elongatus*, *Synechocystis*, and *Anabaena* (6, 18, 19). PconII is a synthetic unregulated core promoter combining -35 and -10 elements of the native *E. coli trp* and *lac* promoters, respectively (20). Here, we developed a set of variant promoters based on PconII that provide a wide range of expression strengths in 5 diverse cyanobacterial strains including a non-model cyanobacterium isolated from an alkaline soda lake. This set of promoters addresses a need for different levels of transcriptional gene expression for metabolic engineering and synthetic biology applications in diverse cyanobacterial strains.

## Materials and Methods

### Strain and Growth Conditions

All strains and plasmids used in these studies are listed in Table S1. Cyanobacterial strains were grown in BG-11 medium as 50-ml or 100-ml liquid cultures in 100-ml or 250-ml flasks, respectively, with orbital shaking (100-125 rpm) or on agar plates (40 ml, 1.5% agar). Culture media were supplemented with appropriate antibiotics as needed: 2 µg mL^−1^ spectinomycin (SPT) plus 2 µg mL^−1^ streptomycin (STR), 5 µg mL^−1^ kanamycin (KAN) and 25 µg mL^-1^ neomycin (NEO). Cultures were grown at 30°C under continuous illumination at 50-100 μmol photons m^-2^ s^-1^, unless otherwise noted. Light measurements were made with a QSL2100 PAR Scalar Irradiance sensor (Biospherical Instruments, San Diego, CA). *E. coli* strains were grown in Lennox broth (LB) liquid medium or on plates with LB medium solidified with 1.2% agar at 37°C with appropriate antibiotics: 100 µg mL−1 ampicillin (AMP), 17 µg mL^−1^ chloramphenicol (CHL), 20 µg mL^−1^ SPT plus 20 µg mL^−1^ STR or 50 µg mL^−1^ SPT, 50 µg mL^−1^ KAN, and 12.5 µg mL^−1^ tetracycline (TET).

### Molecular methods

Plasmid preparations were performed using the QIAprep Spin Miniprep Kit (Qiagen). PCR amplifications were carried out with Q5 High-Fidelity DNA polymerase (New England BioLabs, NEB). Restriction digests followed the supplier’s recommendations (NEB) but with longer incubation times to ensure complete digests. After PCRs or restriction enzyme digestions, DNA purification and concentration were performed using DNA Clean and Concentrator-5 (Zymo Research) and subsequent DNA concentrations were measured using a UV-Vis spectrophotometer NanoDrop 2000c (ThermoFisher). Oligonucleotide phosphorylation was performed using T4 Polynucleotide Kinase (NEB) according to the manufacturer’s instructions, with the modification that PNK reaction buffer and ATP were replaced by T4 DNA ligase reaction buffer. Complementary oligonucleotides were annealed by mixing equimolar amounts (150 to 200 nmol) of each oligonucleotide in a 50 µl reaction containing T4 DNA ligase reaction buffer. The mixture was incubated at 95°C for 5 minutes, then gradually cooled down to 25°C, decreasing the temperature by 1°C every 30 seconds. Assembly of shuttle plasmids were carried out using T4 DNA ligase (NEB) or a GeneArt Seamless Cloning and Assembly Kit (Life Technologies).

### Construction of the PconII* promoter library in S. elongatus

To produce a library of variant PconII promoters, designed as PconII*, with a broad range of promoter strengths, 2 strategies were used. Pairs of oligonucleotides (see Table S1 and Fig. 1A) corresponding to the PconII promoter sequence were synthesized with randomized bases at specific positions in the -10 and extended -10 regions. These oligonucleotides were phosphorylated and annealed to form double-stranded inserts. Two sets of oligonucleotide pairs were designed: set a for strategy 1 and set b for strategy 2. For set a (conII-n5-F_a / conII-n5-R_a, Table S2), the two nucleotides in the extended -10 region and the three most-conserved nucleotides in the -10 region of PconII were randomized. For set b (conII-n5-F_b / conII-n5-R_b, Table S2), the two nucleotides in the extended -10 region and the three least-conserved nucleotides in the -10 region of PconII were randomized. In addition, a pair of oligonucleotides corresponding to the original sequence of the PconII promoter (conII-REF-F / conII-REF-R, Table S2) were used to make a positive control (pAM6017).

**Figure 1.**
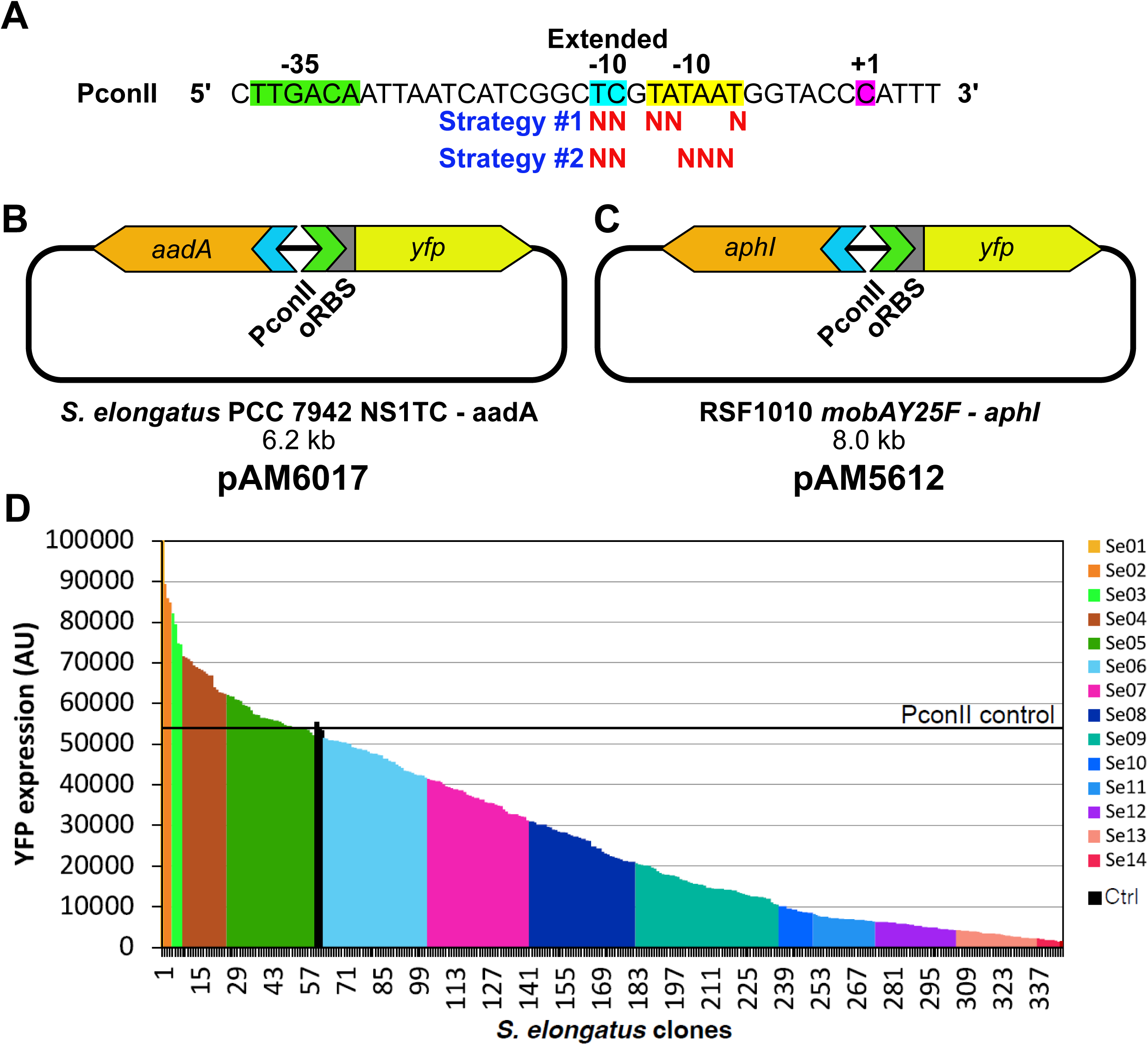
Construction of the PconII* library and assembly of reporter plasmids. (A) PconII sequence with labeled -35 (green highlight), -10 (yellow highlight), and extended -10 regions (blue highlight) and the transcriptional start site at +1 (purple highlight). For each of the 2 strategies to produce variant PconII sequences, “N” is shown below base positions that were randomized. (B) Diagram of *S. elongatus* PCC 7942 neutral-site 1 integration plasmid pAM6017 carrying an *aadA* antibiotic resistance cassette and PconII-YFP reporter gene construct with an optimized ribosomal binding site (oRBS). (C) Diagram of RSF1010-based replicating plasmid pAM5612 carrying an *aphI* antibiotic resistance cassette and PconII-YFP reporter gene construct with oRBS. (D) YFP reporter expression of PconII* library clones in the *S. elongatus* genome. For each promoter variant, YFP expression levels were measured and normalized to Chl*a* measurements. Library clones with similar expression levels were grouped into 14 groups (Se01–Se14) to help follow PconII* clones in subsequent experiments. The expression level of the original PconII promoter is indicated by the black horizontal line.

To generate a library of PconII* promoters driving YFP in *S. elongatus*, we first assembled plasmid pAM5203 to integrate the PconII* library into the chromosome. Plasmid pAM5203 contains 3 devices including: 1, homology sequences for recombination into the *S. elongatus* chromosome at NS1 and an origin of replication for *E. coli*, 2, a SPT and STR resistance gene, and 3, a functional module composed of the *ccdB* toxic gene followed by a ribozyme insulator sequence (RiboJ), a ribosomal binding site (RBS), and a *yfp* reporter gene. Each of these devices were released from CYANO-VECTOR donor plasmids (pAM4818, pAM4836, and pAM5105) and assembled by seamless cloning as described previously (6). Next, pAM5203 was digested with AatII and SwaI to remove the *ccdB* toxic gene and the RiboJ sequence located upstream of an RBS, and the YFP coding sequence. Finally, variant PconII* promoters with an overlapping sequence for the AatII site and a blunt end compatible with the SwaI site were ligated into the linearized pAM5203 backbone to produce the PconII* libraries.

The cloning of variant PconII* promoters for strategy 1 was repeated twice and about 500 *E. coli* colonies were collected and pooled for each cloning experiment. In addition, approximately 1000 colonies were collected from cloning PconII* promoters from strategy 2. The pooled colonies were resuspended in LB medium and plasmid DNA was miniprepped (NucleoSpin Plasmid, Macherey-Nagel) for transformation into *S. elongatus*. For YFP reporter expression experiments, plasmid pAM6017 carrying the original PconII was used as a positive control and pAM5329 carrying the *aadA* resistance gene only was used as negative control.

Transformation of *S. elongatus* followed standard protocols (21) with slight modifications described previously (22). Isolated colonies were grown as small patches on BG-11 plates, then transferred into 2 ml of liquid culture medium and grown in 96-deep-well plates. To determine YFP expression levels, the cultures were adjusted to an optical density at 750 nm (OD750) of 0.1 and YFP fluorescence was then normalized to autofluorescence from chlorophyll *a* (Chl*a*) (Fig. S1). For the PconII* strategy 1 library, we analyzed 626 *S. elongatus* clones after initial cloning experiments and another 311 colonies after an additional cloning experiment. We found only a small number of clones that expressed YFP at levels higher than the PconII control and a large number of clones that expressed YFP at low levels. We selected 68 clones with different expressions strengths from this strategy, but they had only a limited distribution of strong and moderate promoter strengths. These results showed that changing the more conserved nucleotides in the -10 region produced mostly weaker promoters. We therefore designed strategy 2 with the goal of obtaining more strong and medium strength promoter variants. For the PconII* strategy 2 library, 677 clones were grown and analyzed, and we obtained a more even distribution of YFP expression levels resulting in the selection of an additional 275 colonies. A total of 341 colonies were selected from both library construction strategies, and these colonies had a wide distribution of YFP expression levels.

### Construction and characterization of PconII* promoter libraries in diverse strains of cyanobacteria using a broad host range plasmid

Our goal was to characterize the variant PconII* promoters in additional cyanobacterial strains to obtain a small set of promoters that produce a wide range of expression levels in diverse cyanobacteria. The final selected set of PconII* promoters and YFP coding sequences from the integrative *S. elongatus* neutral site 1 plasmids were cloned into a broad-host-range RSF1010-based plasmid (pAM5068), and the resulting plasmids (pAM5068-PconII*-*yfp*) were transferred sequentially into several strains of cyanobacteria as described below.

First, the 341 clones characterized in *S. elongatus* were pooled into 14 groups based on similar YFP fluorescence levels covering ± 5,000 or ± 10,000 AU (Fig. S2). DNA was extracted from each pool, and both the PconII* promoter and the YFP coding sequence were PCR-amplified with primers PLIB-01F/PLIB-01R (Table S2). These PCR products were cloned into the pAM5068 backbone via seamless assembly, following removal of the toxic *ccdB* gene by restriction digestion with SwaI.

For each of the 14 cloning reactions, over 100 *E. coli* colonies were pooled, resulting in 14 small libraries of *E. coli* carrying pAM5068-PconII*-YFP. Each of these plasmid pools were miniprepped and electroporated into an *E. coli* conjugal strain, AM6013. To counter-select against growth of the *E. coli* donor strains following conjugal transfers of the cargo plasmids into cyanobacteria, we used a diaminopimelic acid (DAP) auxotrophic strain of *E. coli*, AM6013, that was generated by introducing the helper plasmid pRL623 carrying MobColK, M.AvaI, M.Eco47II, and M.EcoT22I, and conferring CHL^R^ (23) into WM3064. WM3064, originates from William Metcalf’s laboratory (University of Illinois) and derives from B2155 that carries the RP4 conjugal transfer machinery integrated into its chromosome and is a DAP auxotroph (24, 25).

The pooled libraries were used to transfer the pAM5068-PconII*-YFP plasmids into *Synechocystis* PCC 6803 (hereafter *Synechocystis*) via conjugation. Conjugation methods were performed as previously described (22, 23) with the modification that both *E. coli* donor cultures and conjugation plates (BG-11 with 5% LB) were supplemented with 60 µg/mL DAP (∼315 µM).

A total of 786 *Synechocystis* exconjugant colonies were individually grown, adjusted to an OD_750_ of 0.1, and the YFP fluorescence was measured. Based on this pre-screening, 199 clones were selected with the goal of obtaining a relatively even distribution of YFP fluorescence across all *Synechocystis* clones while also keeping representative clones from each original group in *S. elongatus* to achieve the goal of obtaining a final set of promoters with a wide distribution of expression strengths in diverse cyanobacterial strains. In general, we kept most clones expressing high levels of YFP, greater than PconII reference level, and selected about 1 out of every 5 clones for lower expression levels. Clones with fluorescence levels below that of the negative control were discarded. These 199 clones were re-evaluated, and 24 groups were defined in *Synechocystis* that preserved representation of the original *S. elongatus* groups (Fig. S2).

These clones were then characterized in *Leptolyngbya* BL0902 (hereafter *Leptolyngbya)* (26, 27). Clones from each *Synechocystis* group were pooled, plasmids were extracted, and the new pooled library groups were transformed into the *E. coli* conjugal strain AM6013 and used to transfer the PconII*-YFP plasmids into *Leptolyngbya* via conjugation. A total of 599 *Leptolyngbya* colonies were individually grown, adjusted to an OD_750_ of 0.1, and screened for YFP fluorescence. 112 clones were selected to obtain an even distribution of YFP fluorescence while keeping representative clones from the prior groupings. These were re-evaluated, and 26 groups were defined in *Leptolyngbya* (Fig. S2).

Finally, the resulting clones were characterized in *Anabaena*. Clones from each *Leptolyngbya* group were pooled, plasmids were extracted and transformed into the *E. coli* conjugal strain AM6013. The resulting pooled libraries were used to transfer the PconII*-YFP plasmids into *Anabaena* by conjugation. A total of 262 *Anabaena* colonies were grown, adjusted to OD₇₅₀ = 0.1, and screened for YFP fluorescence. 40 clones were selected to obtain an even distribution of YFP-fluorescence across clones and representation from all previous groupings. Those clones were re-evaluated for YFP fluorescence (Fig. S2), which confirmed the expected expression levels. For the positive control we used plasmid pAM5264 (pAM5068-PconII-yfp), which carries the original PconII promoter driving yfp, and for the negative control we used plasmid pAM5263, which lacks a promoter and yfp reporter gene.

The plasmid DNA from each *Anabaena* clone was extracted, transformed in *E. coli* strain DH5α, and the PconII* sequence was determined. The sequencing results revealed that 15 sequences were duplicates leaving a final set of 25 PconII* promoters for further characterizations.

### Bi-parental conjugation

We performed biparental conjugation using *E. coli* conjugal donor strains to transfer RSF1010-based plasmids into cyanobacterial recipient strains. Electrocompetent DH10B *E. coli* cells carrying the conjugal plasmid pRL443 and helper plasmid pRL623 (23) were electroporated with 10 ng of plasmid DNA and incubated in SOC medium at 37°C for 1 hour. Cells were then plated on LB plates containing CHL (17 µg/mL), AMP (100 µg/mL), and KAN (50 µg/mL). The plates were incubated at 37°C overnight. *E. coli* colonies were selected and grown in LB liquid media with KAN (50 µg/mL) at 37°C overnight. In preparation for conjugation, filamentous cyanobacterial strains (*Leptolyngbya* and *Anabaena*) were fragmented to shorter filaments using probe sonication (20% amplitude for 50 sec., 5 sec. on/off).

Cultures were pelleted by centrifugation and allowed to recover in fresh liquid BG-11 overnight. 1.8 mL of each *E. coli* culture was pelleted by centrifugation, washed twice with LB medium to remove antibiotics, and resuspended in 200 μL LB medium. 30 mL of actively dividing cyanobacterial cultures were pelleted by centrifugation, washed once with BG-11, and resuspended in 6 mL of BG-11. 40 μL of each *E. coli* culture was mixed with 200 μL of each cyanobacterial culture. Each mixture of *E. coli* and cyanobacterial cells was then pelleted by centrifugation at 4000 g for 5 minutes at room temperature and resuspended in 200 μL BG-11. Mixtures were then plated onto Petri plates containing 40 mL BG-11+5% LB solidified with 1.5% Bacto agar. Plates were incubated at 30°C overnight in low light (10-30 μmol photons m^-2^s^-1^) to allow conjugation to occur. The next day, plates were underlaid with 400 µL KAN 100x solution (final concentration 5 µg/mL). Because *Anabaena* is naturally resistant to KAN, NEO (final concentration 25 µg/mL) was instead used to underlay plates containing *Anabaena* exconjugants. All plates were transferred to 70-100 μmol photons m^−2^ s^−1^ illumination at 30°C. After 7-10 days, colonies were selected and streaked onto BG-11 agar plates supplemented with KAN or NEO.

### Fluorescence measurements

Liquid cultures were grown in triplicate for each cyanobacterial strain. Cultures were adjusted to an initial OD750 of 0.1, grown for three days under standard conditions, and adjusted again to an OD750 of 0.1 prior to measurement. A Tecan Infinite® M200 plate reader (TECAN) was used to measure optical densities and fluorescence intensities from 200 μL of culture in black-walled, clear-bottom 96-well plates (Greiner). Excitation and emission wavelengths were set to 490 and 535 nm, respectively, for YFP and 425 and 680 nm, respectively, for Chl*a*.

### Strain Isolation from Environmental Samples

Bioprospected strains ML2A, ML2C1, ML2C2, and ML3B were obtained from enrichment cultures of samples collected from Mono Lake, California. Samples comprised of water and sediment were collected from the lakeshore and used to inoculate liquid BG-11 medium enrichment cultures that were grown with orbital shaking at 30°C with 70-100 μmol photons m^−2^ s^−1^ illumination for 10-14 days. Biparental mating using cells from the mixed enrichment cultures was performed as described above, with the following changes: 200 μL *E. coli* donor cells (instead of 40 μL) and 1 mL mixed culture of cyanobacteria (instead of 200 μL) were used. Environmental samples were sonicated prior to biparental mating to break up cell clumps. 100 μL of the conjugation mixture were plated on BG-11 containing 5% LB and 0.5 M NaHCO_3_ solidified with 1% Gelrite (gellan gum). Plates were incubated at 30°C with 10-30 μmol photons m^−2^ s^−1^ illumination for 24 hours and then transferred to increased illumination of 70-100 μmol photons m^−2^ s^−1^ for 7-10 days. Green colonies of potentially exconjugant cyanobacteria were streaked onto BG-11 plates containing SPT (2 µg/mL) and STR (2 µg/mL) to isolate antibiotic-resistant colonies. To obtain axenic cultures of exconjugant cyanobacterial clones, colonies were re-streaked alternately between BG-11 containing 0.5 M NaHCO_3_ and BG-11 containing SPT and STR. We alternated selection conditions to ensure that high carbonate levels and antibiotic selection were maintained, but the antibiotics were not used on bicarbonate plates due to concerns that high bicarbonate levels can modulate antibiotic susceptibility (28).

Axenic cultures of each cyanobacterial strain were grown in 50 mL of BG-11 in 125 mL flasks under antibiotic selection at 30°C with 70-100 μmol photons m^−2^ s^−1^ illumination. Colony PCRs were performed to amplify the 16S rRNA gene from each strain using cyanobacteria-specific primers 16S27F and 23S30R (29). Sequencing of the 16S rRNA region was conducted using primers 16S378F and 16S784R (29).

To cure strain ML3B of RSF1010-based plasmid pAM5409 we used our previously described protocol (30). Briefly, a culture of ML3B carrying pAM5409 was grown in BG-11 with 0.5 M NaHCO_3_ at 30°C with 70-100 μmol photons m^−2^ s^−1^ illumination but without antibiotics. The culture was passaged regularly to fresh medium when cells reached high density (OD750 > 0.7). After 30 days, culture dilutions were plated on BG-11 agar media to generate single colonies. Single colonies were patched onto both BG-11 agar with and BG-11 agar without antibiotics. Colonies that failed to grow on antibiotic plates indicated loss of plasmid; corresponding patches from non-selective plates were tested for loss of pAM5409. Plasmid loss was confirmed by fluorescence microscopy, which show that the plasmid-cured strain displayed no detectable YFP expression, and by PCR using RSF1010-specific primers (30).

### Construction of Phylogenetic Tree

The phylogenetic tree was constructed based on 16S rRNA gene sequences for identification of strain ML3B (Fig. S4). The evolutionary history was inferred by using the Maximum Likelihood method and General Time Reversible model. The tree with the highest log likelihood (-14053.35) is shown. The percentage of trees in which the associated taxa clustered together is shown next to the branches. Initial tree(s) for the heuristic search were obtained automatically by applying Neighbor-Join and BioNJ algorithms to a matrix of pairwise distances estimated using the Maximum Composite Likelihood (MCL) approach and then selecting the topology with superior log likelihood value. A discrete Gamma distribution was used to model evolutionary rate differences among sites (5 categories (+G, parameter = 0.6326)). The rate variation model allowed for some sites to be evolutionarily invariable ([+I], 56.68% sites). The tree is drawn to scale, with branch lengths measured in the number of substitutions per site. The analysis involved 70 nucleotide sequences with a total of 1362 positions in the final dataset. Evolutionary analyses were conducted in MEGA X. Strain ML3B is marked with a red asterisk (*) and the four standard research strains are marked with red dots (•).

### Microscopy

Micrographs were obtained with a Delta Vision (Applied Precision, Inc.) Olympus IX71 inverted microscope using a 100x oil immersion objective. Images exhibiting cyanobacterial autofluorescence were acquired using TRITC filters (555/28-nm excitation and 617/73-nm emission). Images of yellow fluorescence protein (YFP) were acquired using YFP filters (500/20-nm excitation, 535/30-nm emission). Micrographs were obtained using Resolve3D softWoRx-Acquire v. 4.0.0 and processed using ImageJ (National Institute of Health).

### Strain Cultivation in High Salt

Five cyanobacterial strains were tested: *S. elongatus*, *Synechocystis*, *Anabaena*, *Leptolyngbya*, and Cf. *Nodosilinea* ML3B. All cultures were pre-cultured in BG-11 liquid medium for 4-5 days prior to the experiment. BG-11 buffered with 5 mM MOPS was prepared with different concentrations of sodium bicarbonate (NaHCO_3_): no addition, 0.1 M, 0.25 M, 0.5 M, 1 M, and 2 M. Each strain was added to 2-mL cultures of BG-11 at each concentration of NaHCO_3_ and adjusted to an OD750 of 0.1. All culture assays were performed with biological triplicates using clear 6-well plates (Greiner). Cultures were grown for 5 days, with orbital shaking at 30°C under continuous illumination with 60-80 μmol photons m^-2^ s^-1^. OD750 readings for cell density were measured daily using a Tecan Infinite® M200 plate reader.

## Results & Discussion

### Generation of the promoter library

We generated different promoter sequences by randomizing nucleotides in the -10 and extended - 10 regions of PconII. The variant promoters (designated PconII*) were produced using oligonucleotide pairs with randomized bases at specific positions that were annealed with each other and cloned upstream of a reporter gene. To obtain a wide range of promoter strengths, we used two strategies that targeted the 2 nucleotides in the extended -10 region and either the 3 most-conserved or the 3 least-conserved bases in the -10 region of PconII (Fig. 1A). These positions are also conserved across promoters in cyanobacterial strains including *Synechocystis*, *Anabaena*, and *S. elongatus* (31–33).

The library of PconII* promoters was first screened in *S. elongatus*. We used the *yfp*-reporter vector pAM5203 to integrate PconII*-*yfp* reporter constructs into neutral site 1 in the chromosome (Fig. 1B). Chromosome neutral sites in *S. elongatus* allow integration of DNA sequences without affecting normal cell phenotypes (21). Individual clones were randomly selected for measurement of YFP expression, which was normalized to autofluorescence from Chl*a*. Modification of the most conserved nucleotides in the -10 region (“Strategy 1” primers) produced promoters skewed towards high and low YFP expression with poor representation of intermediate expression levels (Fig. S1). Modification of the least conserved nucleotides in the -10 region (“Strategy 2” primers) yielded a more even distribution of expression levels of the PconII* variants (Fig. S1). Clones of PconII* promoters made using both strategies were grouped together as a single library for subsequent experiments. A total of 1643 clones were initially analyzed for YFP expression in *S. elongatus*, and 341 PconII* promoter clones with a wide range of expression levels were selected for further characterization as described in Materials and Methods (Fig. 1D).

### Cloning PconII* promoters in a broad host-range vector and selection of a set of promoters for diverse cyanobacterial strains

To obtain a characterized set of variant PconII* promoters with a wide range of promoter strengths in diverse cyanobacterial strains, subsets of the PconII* promoter library were introduced and evaluated in 3 additional cyanobacterial strains: *Synechocystis*, *Leptolyngbya*, and *Anabaena*. *S. elongatus* and *Synechocystis* are unicellular model cyanobacterial strains. *Anabaena* is a model filamentous strain that produces specialized heterocyst cells for nitrogen fixation (34). *Leptolyngbya* is a filamentous nonheterocystous strain that is tolerant of high salt and high-pH growth conditions and may be suitable as a potential production strain (26, 27). GC content of the strains varied widely, with *Anabaena* at 41.5%, *Synechocystis* at 47.5%, *S. elongatus* at 55.5%, and *Leptolyngbya* at 57.5% (35–38). Together, these organisms represent a phylogenetically diverse range of cyanobacteria (Fig. S4).

Variant PconII* promoters from the initial library in integrative vector pAM5203 were cloned into the self-replicating vector pAM5068. Briefly, the 341 *S. elongatus* clones containing PconII*-yfp were pooled according to promoter strength (YFP expression) into 14 groups (Fig. 1) and genomic DNA was extracted from each group. Then, the PconII* variant sequences driving *yfp* from each pool were PCR-amplified and cloned into the broad-host-range self-replicating RSF1010-based vector pAM5068 (39). Plasmid pAM5068 carries a KAN resistance gene (*aphI*) and its own replication machinery that allows its transmission and maintenance in diverse strains of cyanobacteria (30, 39) (Fig. 1C). The resulting plasmids (pAM5068-PconII*-yfp) were sequentially transferred by conjugation into the 3 additional strains of cyanobacteria with the goal of producing a single set of expression plasmids with a wide distribution of promoter strengths in each of the 4 different strains. The 14 groups of pAM5068-PconII*-yfp plasmids, which represented the 341 different promoters initially characterized in *S. elongatus*, were transferred into *Synechocystis* and isolated colonies were individually grown, evaluated for YFP expression and grouped by promoter strength into 24 new groups. The promoters in these groups were then similarly characterized in *Leptolyngbya* and then finally in *Anabaena* (see Materials and Methods for details). During the characterization of promoter strengths in each strain, clones were selected to ensure representation of the range of promoter strengths in the other strains. Screening the pAM5068-PconII*-yfp library in all 4 cyanobacterial strains resulted in a set of 40 library clones (Materials and Methods and Fig. S2).

### Characterization of a subset of variant PconII* promoters with a wide range of expression strengths in diverse strains

The PconII* sequences for the 40 clones were determined, and after removing duplicate sequences, we obtained a final set of 25 plasmids in vector pAM5068 with PconII* promoters of graded strengths driving a *yfp* reporter (Supplement File S1). For subsequent experiments, we included a positive control construct with the original PconII promoter (pAM5612) and a negative control construct lacking a promoter (pAM5729). Hereafter, the characterization of the variant PconII* promoters in all strains, including in *S. elongatus,* were conducted using these 27 broad-host-range RSF1010-based plasmids.

We assessed the expression strengths of the final set of 25 variant PconII* promoters plus 2 controls in the 4 model strains. The broad host-range plasmids were introduced individually by natural transformation into *S. elongatus*, and by conjugation into *Synechocystis*, *Anabaena*, and *Leptolyngbya*. Because RSF1010-based plasmids are not stably maintained in *S. elongatus*, we used a Δ*ago* mutant of *S. elongatus*, strain AMC2664, that allows the replication and maintenance of RSF1010-based plasmids (22). Characterization of the promoters in each of the four strains produced a wide range of reporter expression levels. In *S. elongatus*, expression ranged from 4% to nearly 125% of the level of the original PconII promoter (Fig. 2A). For these experiments, because the PconII*-*yfp* cassettes are carried on a replicative plasmid rather than integrated into the *S. elongatus* chromosome at a neutral site, there were some differences in YFP expression levels. However, the distribution of plasmid-based promoter strengths remained spread across a wide range of expression levels in *S. elongatus*, similar to the initial characterization of the PconII*-*yfp* constructs integrated into the chromosome (Fig. 1D).

**Figure 2.**
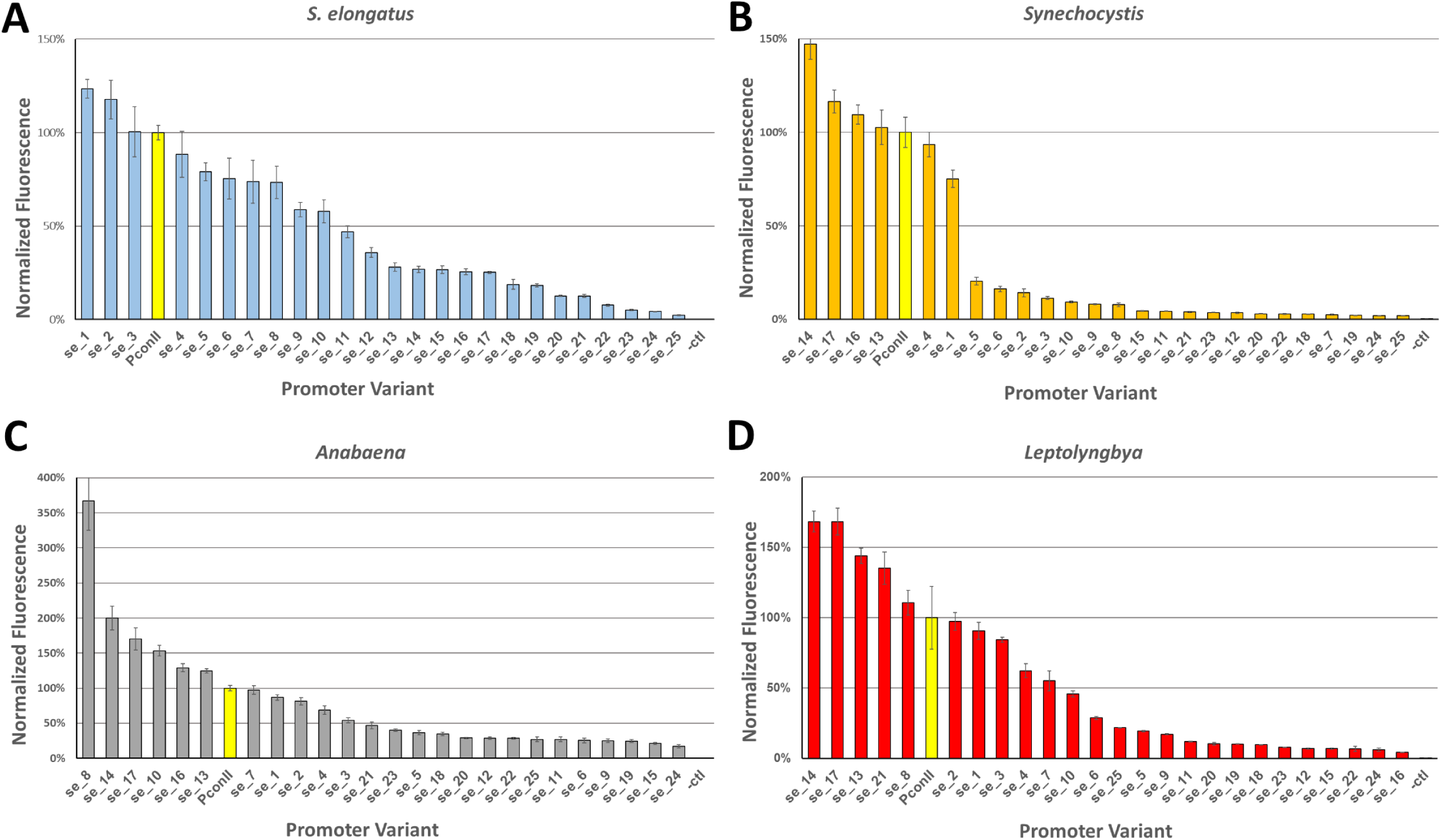
Yellow fluorescent protein (YFP) reporter expression levels of 25 selected PconII* library clones in 4 cyanobacterial strains. Cell cultures with each promoter variant were grown for 5 days in multi-well plates, adjusted to an OD750 of 0.1, and measured for YFP expression. Expression levels were normalized to those of the original PconII promoter (yellow bars). Error bars represent standard error of the mean calculated from biological triplicates.

In the other tested strains, we observed a similarly wide range and distribution of promoter expression levels. In *Synechocystis*, the set of 25 promoters spanned a 150-fold range in expression levels (Fig. 2B). Some promoters with high expression in *S. elongatus* were much lower in *Synechocystis* relative to the unaltered promoter. For example, variant PconII*_se_1 is the highest-expressing promoter in *S. elongatus,* at 123% of PconII, but the same variant only exhibits 75% of the PconII expression level in *Synechocystis*. In contrast, the highest-expressing promoter in *Synechocystis* was PconII*_se_14 at 147% of PconII but has only 26% of PconII activity in *S. elongatus*. In *Anabaena*, most of the PconII* variants span a 200-fold range in expression levels, although one promoter, PconII*_se_8, had an expression level 3.6 times higher than PconII (Fig. 2C). In *Leptolyngbya*, we observed a 170-fold range of promoter strengths (Fig. 2D). We note that many of the variant PconII* promoters had expression strengths in *S. elongatus* that differed from the expression strengths in the other three strains (Fig. 2, Supplemental File S1). For example, variants PconII*_se_14 and PconII*_se_17 were consistently high-expressing promoters across *Synechocystis*, *Anabaena*, and *Leptolyngbya*, but had relatively low expression strengths in *S. elongatus*.

We selected three PconII* variants with low, medium, and high YFP expression levels in each strain to assess promoter expression using fluorescence microscopy. These micrographs confirm the previous ranking of promoter strengths, with weaker PconII* variants exhibiting the lowest fluorescence intensity and stronger variants exhibiting the highest (Fig. 3A-D).

**Figure 3.**
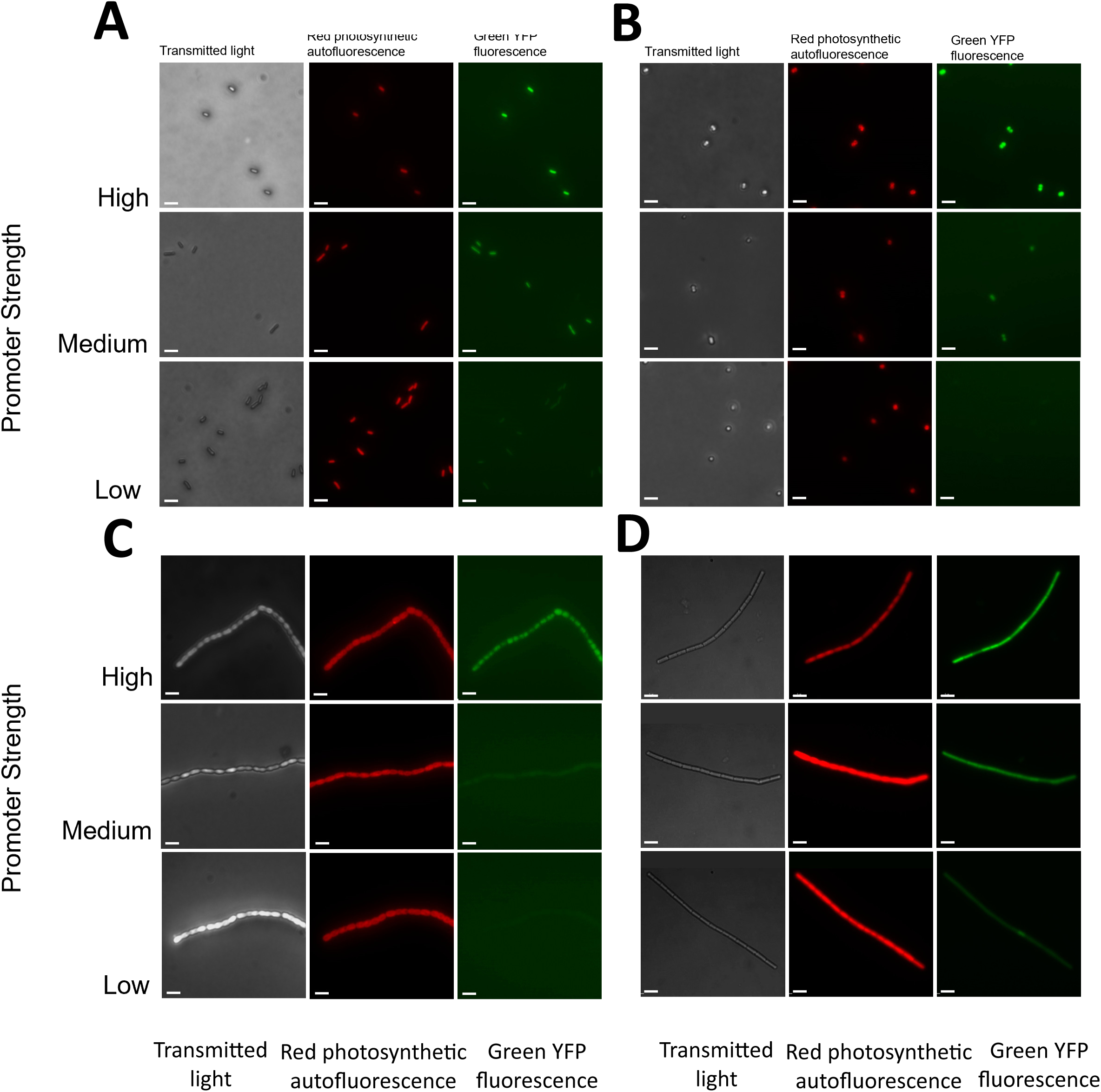
Micrographs of cyanobacterial strains expressing low, medium, and high strength PconII* variants driving a *yfp* reporter gene. (A) *S. elongatus*, (B) *Synechocystis*, (C) *Anabaena*, and (D) *Leptolyngbya*. For each strain: left panels, transmitted light; middle panels, red autofluorescence from photosynthetic pigments; right panels, green YFP fluorescence. Scale bars, 5 μm. The following promoters were used: *S. elongatus:* low, PconII*_se_21; medium, PconII*_se_10; high, PconII*_se_1. *Synechocystis:* low, PconII*_se_2; medium, PconII*_se_1; high, PconII*_se_14. *Anabaena:* low, PconII*_se_23; medium, PconII*_se_17; high, PconII*_se_8. *Leptolyngbya*: low, PconII*_se_9; medium, PconII*_se_3; high, PconII*_se_14.

The wide range of expression strengths in the final set of 25 promoters should be useful for many applications. For example, in a previous study that engineered *S. elongatus* to produce ethylene (40), metabolic intermediates in the tricarboxylic acid (TCA) cycle were depleted, which led to a decline in cell growth. For similar metabolic engineering, the set of PconII* promoters would be useful to fine-tuned metabolic flux of production pathways to ensure replenishment of intermediate compounds and decrease production of toxic intermediates to achieve optimized production of final products.

The set of PconII* promoters in the RSF1010-based broad host-range vector should be useful for a number of applications in diverse strains. However, the promoters may need to be moved into other vectors for some applications. For example, similar to *S. elongatus*, some strains may not be compatible with the RSF1010-based plasmids used in this study. We addressed this limitation by using the engineered AMC2664 Δ*ago* strain of *S. elongatus* (22). Our characterization of this strain indicates that it has a normal phenotype and that it should be suitable for research or other applications. The RSF1010 replicon has the advantage of replicating in diverse bacterial hosts but its copy number may vary depending on the strain and growth conditions (10, 41) thereby affecting expression levels. In addition, because promoters can have different expression levels in different sequence contexts and in different strains, promoter strengths need to be validated for each specific application.

### Characterization of the promoter set in a newly isolated strain

To demonstrate the utility of the set of PconII* promoters, we characterized them in a newly isolated genetically tractable cyanobacterial strain. Plasmids based on the broad-host-range RSF1010 plasmid can serve both as a tool for conducting genetic modifications in model cyanobacteria and as a practical method for isolating new genetically tractable strains through bioprospecting, which is the discovery of new organisms in natural habitats for biotechnological or other applications (30). In this study, we screened for environmental strains that were halotolerant and capable of growth in high levels of bicarbonate. Halotolerant strains are useful due to the appeal of using waste or brackish water for sustainable large-scale cultivation of cyanobacteria for biotechnology applications, and high-bicarbonate media can increase growth rate and carbon fixation (42–44).

Water samples with sediments were collected from the shore of Mono Lake, CA, an alkaline soda lake, and used to grow mixed algal cultures in modified BG-11 medium under constant light at 30°C. The broad host-range YFP reporter plasmid pAM5409 was conjugated into the mixed cultures, as previously described (30). After antibiotic selection, colonies were picked and grown in individual liquid cultures and then observed by microscopy to distinguish between cyanobacteria and eukaryotic green algae based on cell size and morphology. We isolated four different genetically tractable cyanobacterial strains with distinct 16S rRNA gene sequences (Table S3). The four exconjugant strains were grown on selective media and each demonstrated YFP fluorescence (Fig. S3). All four strains were filamentous and unbranched, with three forming straight filaments, and one strain, ML2C1, forming helical filaments (Fig. S3). All four strains formed isolated colonies on agar plates. Three of the four strains routinely flocculated, forming cell clumps when grown as liquid cultures in flasks. One strain, ML3B, grew planktonically without significant clumping.

We selected strain ML3B for additional characterization and genetic experiments because it grows as dispersed filaments in liquid medium which makes culture optical-density and fluorescence measurements more accurate. Non-clumping strains also facilitate maintenance of axenic cultures (45). To obtain the ML3B strain cured of the pAM5409 reporter plasmid used for the initial isolation, we grew the exconjugant ML3B strain in liquid growth medium without antibiotics and passaged the culture every 3-4 days for 30 days. Single colonies were tested for loss of the plasmid by screening for antibiotic susceptibility. The resultant cured strain was susceptible to antibiotic selection and did not express YFP fluorescence. The loss of the pAM5409 plasmid was confirmed by PCR (data not shown). Morphologically, ML3B grows as non-branching filaments, with cells measuring 1.17 ± 0.12 μm long and 1.16 ± 0.13 μm wide. Phylogenetic analysis suggested that ML3B belongs to the *Nodosilinea* genus (Fig. S4). ML3B is naturally resistant to KAN, but susceptible to NEO, SPT, and STR at the same concentrations that we use for other cyanobacterial strains (see Materials and Methods). It does not form heterocysts and did not grow on medium lacking a source of fixed nitrogen. ML3B was tolerant of high-bicarbonate media and grew well at a bicarbonate concentration of 0.5 M, which none of the four laboratory strains can survive (Fig. S5).

We conjugated the set of reporter plasmids with the variant PconII* promoters into the cured ML3B strain to characterize the promoter strengths in this new strain. The PconII* promoters produced a 150-fold range of expression levels with a relatively smooth distribution of strengths in ML3B (Fig. 4A). We selected 3 strains with low, medium, and high strength PconII* promoters to image via fluorescence microscopy. As expected, levels of YFP fluorescence observed by microscopy paralleled those measured from cultures using a plate reader (Fig. 4B).

**Figure 4.**
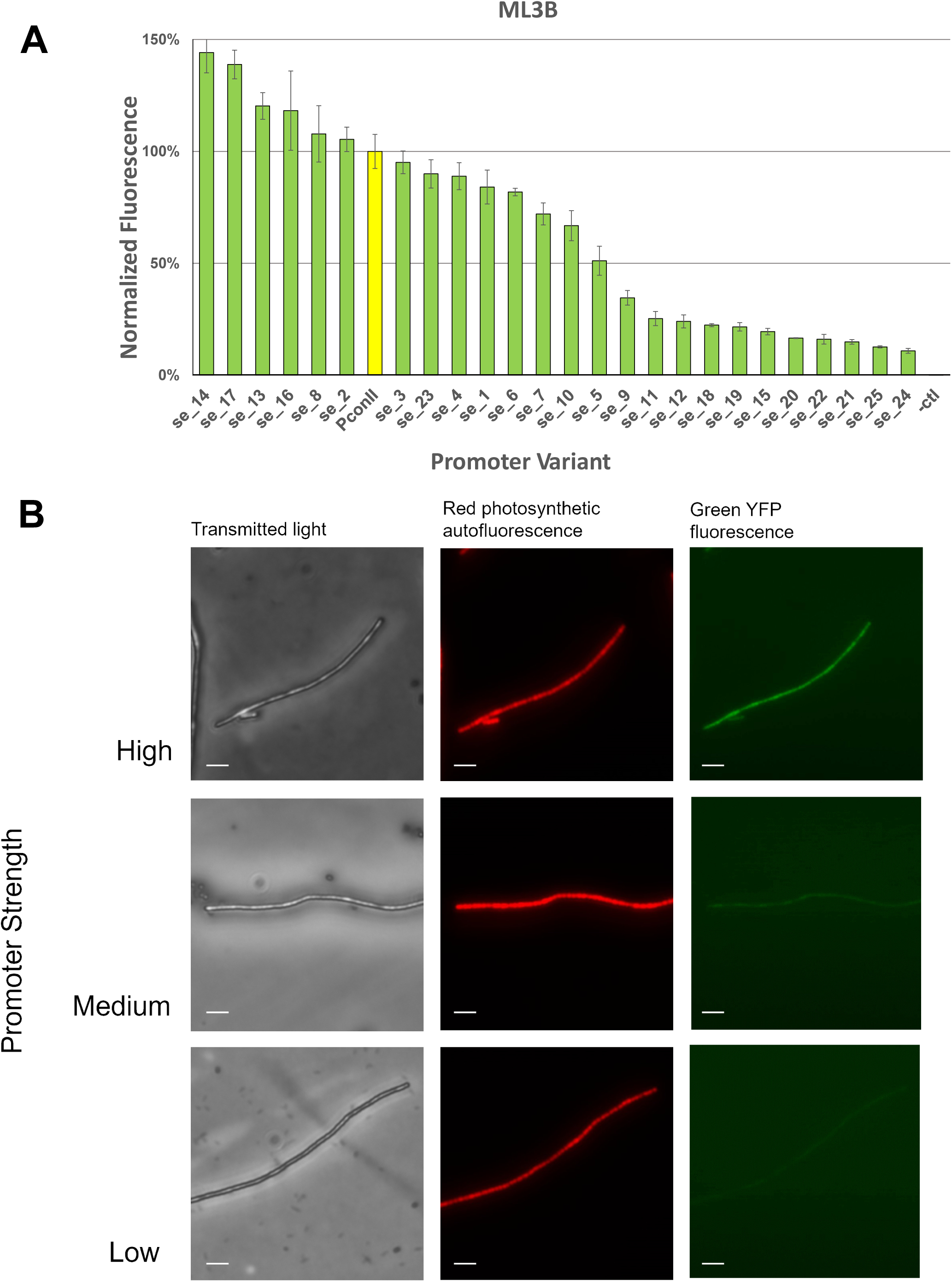
Characterization of the set of PconII* promoter variants in bioprospected strain ML3B. (A) Fluorescence levels of ML3B strains with the PconII* promoter variants. Cell cultures with each promoter variant were grown for 5 days in multi-well plates, adjusted to an OD750 of 0.1, and measured for YFP fluorescence. Expression levels are normalized to the expression level of the PconII promoter (yellow bar). Error bars represent standard error of the means from biological triplicates. (B) Micrographs of strain ML3B expressing low, medium, or high strength PconII* variants driving *yfp*. Left panels, transmitted light; middle panels, red autofluorescence from photosynthetic pigments; right panels, green YFP fluorescence. Scale bars, 5 μm. The following promoters were used for ML3B: low, PconII*_se_24; medium, PconII*_se_15; high, PconII*_se_4.

These experiments demonstrated that strain ML3B, an extremophile cyanobacterium capable of growth at high salinity and alkalinity, is suitable for genetic manipulation, and that the subset of 25 PconII* promoters exhibited a wide range of promoter strengths in this newly isolated strain.

Identifying new genetically tractable cyanobacterial strains from different environments is useful for both basic scientific research and biotechnology applications. For example, strains that can be grown to high densities in non-potable brackish water would allow sustainable large-scale production of biomass and desired products without competing for freshwater resources required for food production and human consumption. Cultivation of algae has been proposed as a strategy for removing excess nutrients and elevated levels of dissolved carbonate salts from agricultural wastewater while producing valuable biomass and natural products (46, 47). Additionally, harsh growth conditions can limit contamination from undesirable strains and predators in large-scale growth facilities (26).

Bioprospecting methods designed for identifying genetically tractable cyanobacteria enable the identification of new strains for biotechnology applications and large-scale industrial growth (30). In this study, we isolated four genetically tractable cyanobacterial strains from an alkaline soda lake with high alkalinity and dissolved carbonate levels, making them promising candidates for cultivation in brackish non-potable water. One of the strains, ML3B, was selected for further characterization. Phylogenetic analysis showed that ML3B is closely related to a group of cyanobacteria isolated from hypersaline, alkaline lakes in western Brazil. ML3B is most closely related to strain *Nodosilinea* sp. CENA 523 (48) (Fig. S4). Another phylogenetically related strain is *Leptolyngbya* sp. KIOST-1 isolated from a culture pond of *Arthrospira* in South Korea; this salt-tolerant strain, with high protein content comprising more than 50% of cellular biomass, was proposed to be a potential biomass producer comparable to *Arthrospira* (49) (Fig. S4). Based on its characteristics and those of its phylogenetic neighbors, strain ML3B may possess the qualities that make it suitable for large-scale cultivation at bicarbonate levels as high as 0.5 M (Fig. S5). ML3B could also serve as a genetic model strain for studying the metabolic capabilities that allow it to survive in hypersaline and alkaline conditions.

### Sequence comparison of promoter variants

We examined the sequences of each of the final 25 PconII* variants in the 4 model strains as well as the newly isolated strain Cf. *Nodosilinea* ML3B (Fig. 5). To determine if particular nucleotide changes correspond to patterns of the higher or lower relative expression across the tested strains, we analyzed the sequences of each of the PconII* variants. PconII* promoters with a stronger expression than PconII often only had changes in the extended -10 region at positions -15 and -14. The overall strongest promoter was PconII*_se_14, which had changed only the original “TC” at positions -15 and -14 to “CT”. Changes in the least conserved bases of the -10 region, the “TAA” at positions -10 to -8, almost always yielded promoters weaker than PconII, with a few notable exceptions. For instance, the strongest promoter in *S. elongatus,* PconII*_se_1 has the same “CT” modification at -15 and -14 as PconII*_se_14 but has a “TAA” to “TAC” change at -10 to -8. This change lowered the expression level in all other strains but increased it in *S. elongatus.* As noted previously, *S. elongatus* was somewhat of an outlier in promoter expression strengths compared to other strains, and all of the stronger promoters in *S. elongatus* had changes to the “TAA” at -10 to -8. Only a few promoters had changes at the most conserved positions “TA…T” at -12, -11 and -7, which in most cases resulted in weaker expression than PconII. One exception was PconII*_se_8, which was the strongest promoter in *Anabaena* and ranked as the fifth strongest promoter in *Leptolyngbya* and ML3B. It is also apparent that making more changes to the promoter region generally lowered expression, with the weakest overall promoters having 5 base changes (Fig. 5).

**Figure 5.**
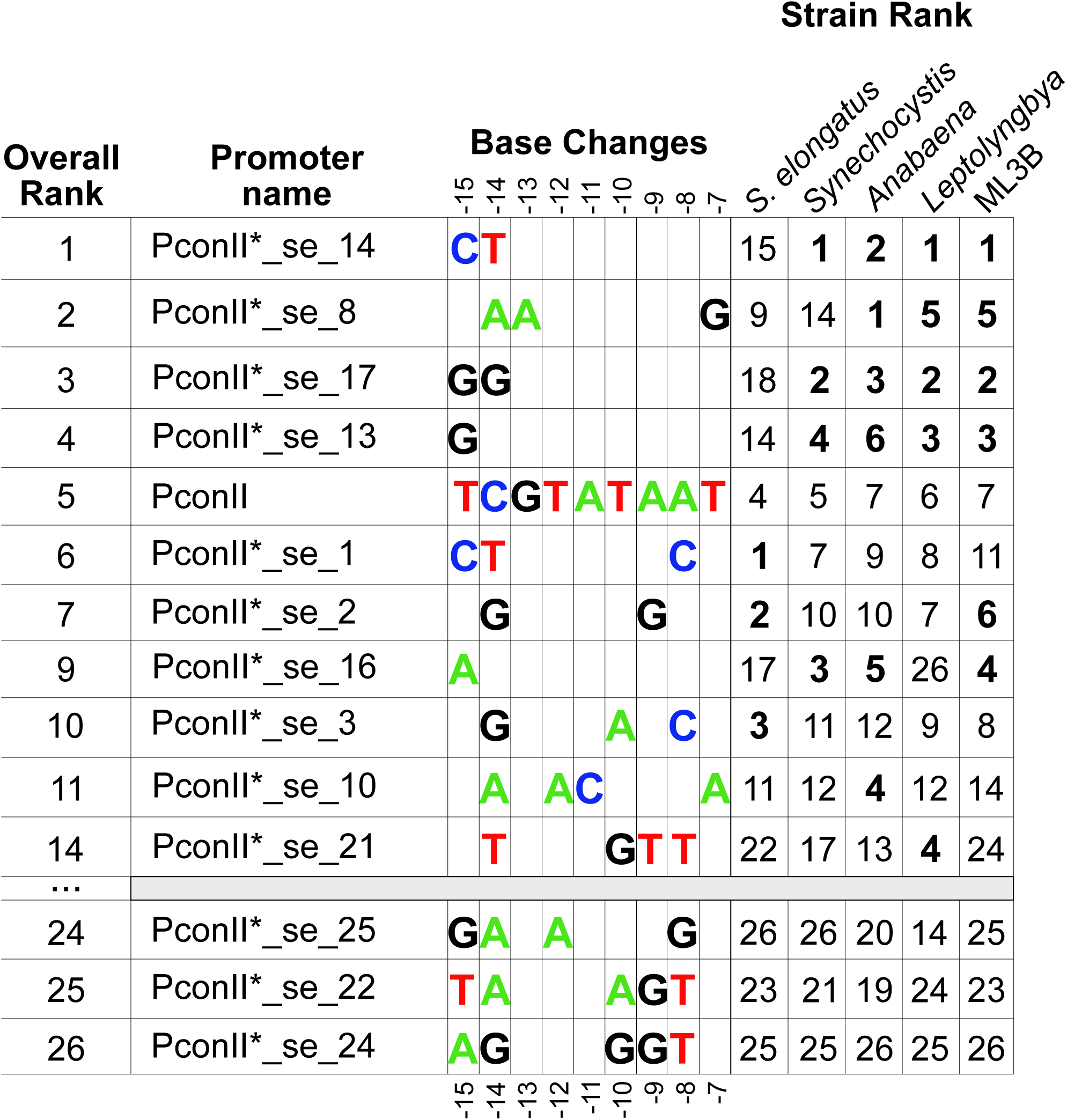
Base changes of PconII* variants with stronger expression than PconII. Library clones were characterized in five cyanobacterial strains by YFP expression level normalized to the original PconII promoter in the same strain (Fig. 2 and 4). Overall ranking of promoter strengths were calculated by averaging normalized expression levels in the five strains, then sorting variants based on decreasing YFP expression strength (Table S1). All variants ranked stronger than PconII in any strain were included, along with three examples of the weakest variants across all strains. The relative rank in each strain is shown; variants stronger than PconII are indicated in bold. The original PconII sequence is shown; positions with unchanged nucleotides in the PconII* variants are not shown. The number of stronger versus weaker promoter variants in each strain were: *S. elongatus* (stronger: n=3, weaker: n=22); *Synechocystis* (stronger: n=4, weaker: n=21); *Anabaena* (stronger: n=6, weaker: n=19); *Leptolyngbya* (stronger: n=5, weaker: n=20); and ML3B (stronger: n=6, weaker: n=19).

In *Synechocystis,* the core promoter sequence appears to be more sensitive to changes because all changes in the -10 core “TATAAT” weakened promoter expression. We also observed a limited number of intermediate expression levels for PconII* promoters in *Synechocystis* (Fig. 2B). This lower representation of PconII* sequences in the middle range of promoter strengths was also seen in the initial characterization of the library in *Synechocystis* (Fig. S2B). Unlike the other strains, the only mutations that increased *Synechocystis* promoter strength were changes to the extended -10 region (Fig. 5).

### Conclusions

In this study, we generated a library of PconII* constitutive promoters that were initially characterized in *S. elongatus* and then evaluated in 3 additional strains of cyanobacteria to create a set of promoters with a wide range of expression levels. This final set of 25 variant PconII* promoters were characterized in each of the 4 diverse model strains and a cyanobacterial strain newly isolated from a saline soda lake. The set of PconII* promoters produced YFP reporter expression levels that ranged from very low, near background levels of a negative control, to levels surpassing the strong expression levels of the original PconII promoter (Fig. 2A-D and Fig. 5). Many native and heterologous promoters have been used in cyanobacteria, but they are often characterized in a single strain (1, 3, 4, 12). The promoters produced in this work were characterized in diverse cyanobacterial strains and shown to produce a broad, graded range of expression levels. Therefore, this set of promoters would be useful for cyanobacterial fundamental research, metabolic engineering, or synthetic biology applications.

## Acknowledgements

We thank Ryan Simkovsky, Jeffrey T. Mindrebo, and Yvone Reyes for their technical assistance during the construction of the initial library of PconII* promoters. This work was supported by the United States Department of Energy, Office of Energy Efficiency and Renewable Energy, under Contract Number EE0008246, the National Institute of General Medical Sciences of the United States National Institutes of Health under Award Number R01GM118815, and the National Science Foundation through the UC San Diego Materials Research Science and Engineering Center (UCSD MRSEC) DMR-2011924. The content is solely the responsibility of the authors and does not necessarily represent the official views of the DOE, NIH, or NSF.

A.T., B.B., and J.W.G. conceptualized the work and designed experiments. J.W.G. provided funding and resources. B.P.T. and K.P.T. performed experiments and analyzed data. B.B. and A.T. supervised research and laboratory experiments. K.P.T. and B.B. wrote the original manuscript. B.B, A.T., and J.W.G. edited the manuscript. All authors read and approved the final manuscript.

